# A simplified disease resistance assay using YFP-expressing *Potato Virus X* in *N. benthamiana* reveals a cell death-independent immune function of RBA1

**DOI:** 10.1101/2023.10.20.563110

**Authors:** Keiichi Hasegawa, Ton Timmers, Jijie Chai, Takaki Maekawa

**Affiliations:** Institute for Biochemistry, University of Cologne, 50674 Cologne, Germany; Max Planck Institute for Plant Breeding Research, 50829 Cologne, Germany; Cluster of Excellence on Plant Sciences (CEPLAS), Cologne, Germany; Institute for Plant Sciences, University of Cologne, Cologne, 50674, Germany; School of Life Sciences, Westlake University, Zhejiang China

## Abstract

R (resistance) proteins, such as intracellular NLRs (nucleotide-binding leucine-rich repeat receptors), are integral components of the plant innate immune system (van Wersch et al., 2020). Host responses following R protein activation include the generation of reactive oxygen species, sustained increases in cytosolic Ca^2+^, transcriptional reprogramming and, typically, rapid host cell death at sites of pathogen infection, which together ultimately lead to pathogen growth restriction (Wang et al., 2023). To assess the activity of R proteins, agroinfiltration-mediated transient gene expression assays have been widely used in *Nicotiana* species (e.g., *N. benthamiana*). In these transient assays, host cell death is often chosen as an indicator of R protein activity from the host responses mentioned above, in part because of the ease of experimentation. However, the extent to which host cell death is a proxy for disease resistance signaling has long been debated, as host cell death and pathogen growth restriction can be uncoupled in several cases (Bendahmane et al., 1999; Coll et al., 2010; Heidrich et al., 2011, Maekawa et al., 2023). To assess the disease resistance activity of R proteins, bacterial growth assays have been employed in combination with transient *R* gene expression in *N. benthamiana* (Sun et al., 2021). Bacterial growth assays, however, require multiple experimental procedures, including agroinfiltration, pathogen infection and bacterial counts, which hinders high-throughput studies of *R* gene-mediated disease resistance. Here, we report a simple plate reader-based assay to assess *R* gene-mediated disease resistance activity against PVX (Potato virus X) that expresses YFP (PVX-YFP). Unlike bacterial pathogens, PVX proliferation in *N. benthamiana* is not restricted by the intrinsic activity of the EDS1 signaling pathway as previously shown by virus-induced *NbEDS1* gene silencing (Peart et al., 2002) and as we consistently show in this study using a *Nbeds1* gene knockout mutant. This feature would increase the sensitivity of the assay, allowing it to capture a weak-to-moderate disease resistance activity of R proteins, as the contribution of basal immunity to PVX via the *Nb*EDS1 pathway is negligible. Using this assay, we show that a non-cell death-inducing mutant of the R protein of RBA1 (Response to HopBA1), which lacks 2′,3′-cAMP/cGMP synthetase activity but retains NADase activity, confers PVX resistance in an EDS1 signaling pathway-dependent manner.

---

PVX is a single-stranded RNA virus and expression of full-length PVX cDNA via agroinfiltration-mediated gene expression allows PVX replication in *Nicotiana* species (Larsen and Curtis, 2012). Insertion of a cDNA of fluorescent protein under a duplicated copy of the viral coat protein promoter allows tracking of PVX replication in plants (Peart et al., 2002). Unlike bacterial growth assays in *Nicotiana* species, in which pathogen infection takes place a few days after agroinfiltration, in the PVX disease resistance assay agroinfiltration and pathogen infection can be performed simultaneously (Collier et al., 2011), with minimal experimental efforts. In this study, we generated a binary vector that allows for PVX replication along with monomeric YFP expression and used YFP intensity as a proxy for PVX viral load. To quantify PVX-produced YFP in *N. benthamiana* leaf extracts, we optimized a fluorescence plate reader-based measurement (Fig. 1A), which is simpler than quantification by western blot. Briefly, in this method agrobacteria carrying the PVX-YFP binary vector and agrobacteria carrying a *R* gene were co-infiltrated into *N. benthamiana* leaves. Four days after infiltration, leaf discs were collected from the infiltrated areas using a biopsy punch. The leaf protein extracts were used to measure YFP intensity (detailed method is described in the Supplemental Methods). First, we determined the emission spectra of leaf extracts containing PVX-YFP or PVX without fluorescent protein (Fig. 1B). The emission spectrum of PVX-produced YFP was in good agreement with the canonical spectrum of YFP in the leaf extract, and PVX replication in leaves did not produce fluorescence that overlapped with the YFP spectrum (Fig. 1B). The YFP intensity of a dilution series using a leaf protein extract from non-transformed leaves showed that this method has a high dynamic range with a linear correlation between fluorescence and dilution factor and that a detergent, Triton X-100, can be omitted from the extraction buffer (Fig. 1C). Furthermore, the quantified YFP fluorescence intensities in the plate reader-based measurement corresponded to some extent with the fluorescence images (Fig. 1D). Consistent with a previous study (Peart et al., 2002), PVX-YFP proliferation was comparable in wild type and the *eds1 pad4 sag101a sag101b* mutant of *N. benthamiana* (Fig. 1E). Co-expression of a wheat NLR, Sr35 and its cognate effector, AvrSr35 (Förderer et al., 2022), caused host cell death in *N. benthamiana* in the presence of PVX-YFP (Fig. 1F), suggesting that agrobacterium-mediated delivery of PVX-YFP T-DNA does not suppress NLR-induced host cell death. Furthermore, the very low YFP intensity (average intensity ± standard error = 1991 ± 402, n=4), which is nearly equivalent to the YFP intensity of leaf protein extract from non-transformed leaves (Fig. 1C), indicates that host cell death does not result in the production of metabolites that show autofluorescence in the YFP assay.

**Figure 1.**
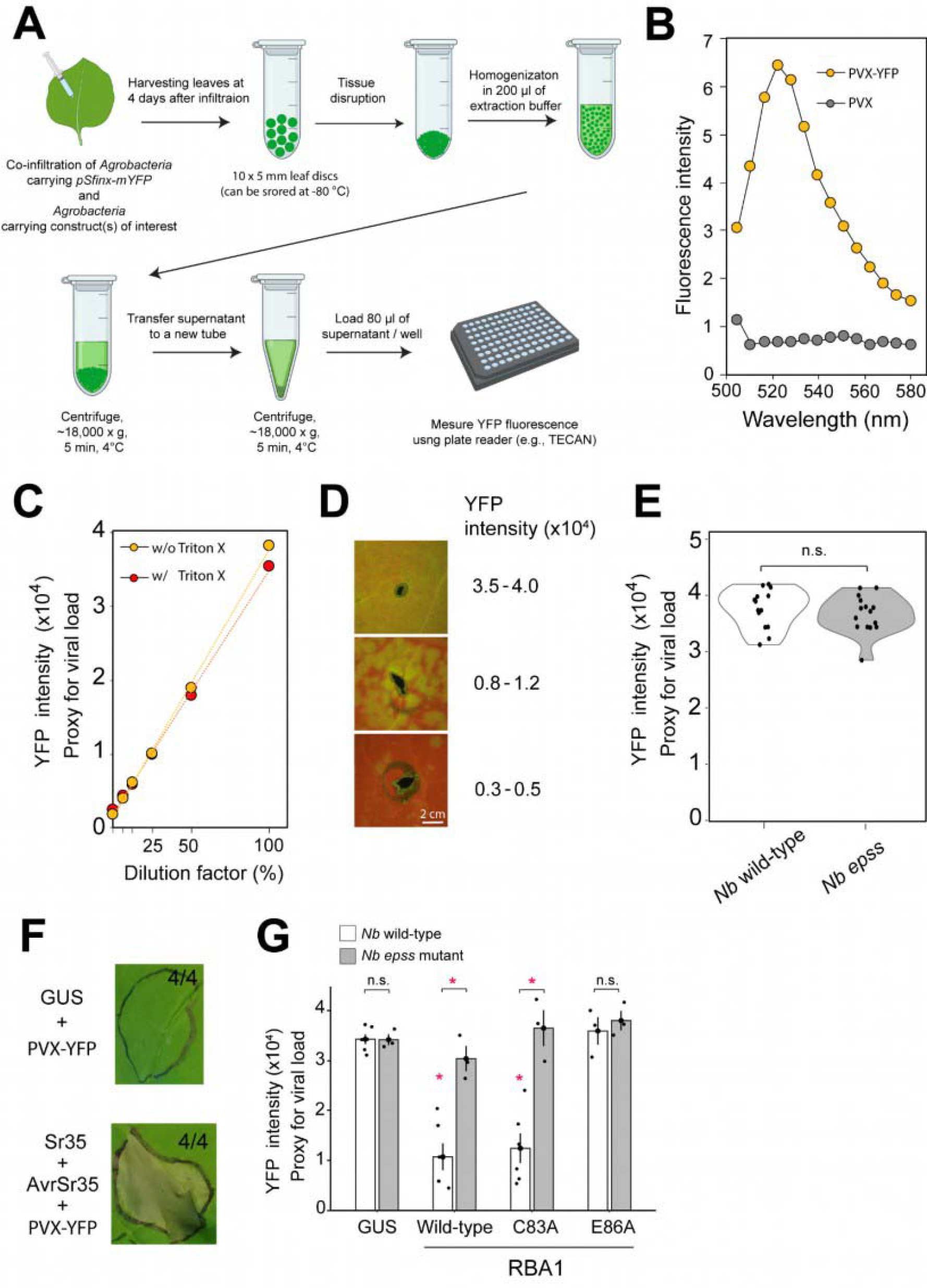
Disease resistance assay using *Potato Virus X* expressing YFP in *N. benthamiana*. **A)** Schematic representation of the disease resistance assay using Potato Virus X expressing YFP (PVX-YFP). **B)** The emission spectra for leaf extracts expressing PVX-YFP or PVX. **C)** A linear correlation between fluorescence and dilution factor in a leaf lysate in the presence and in the absence of 0.1% Triton X-100. Leaf lysates from non-transformed plants were used for dilution. **D)** Representative fluorescence stereomicroscopic images of *N. benthamiana* leaves infiltrated with PVX-YFP with corresponding YFP intensity measured with the TECAN plate reader. The leaves were examined at four days after agroinfiltration. YFP intensity measured with the plate reader from two biologically independent samples is shown. **E)** Violin plots showing the degree of dispersion of the quantified YFP intensities from leaves co-expressing GUS and PVX-YFP in *N. benthamiana*. **F)** Expression of Sr35 and YFP-expressing potato virus X (PVX) caused host cell death *in N. benthamiana* (wild-type). β-glucuronidase (GUS), Sr35, and AvrSr35 (Förderer et al., 2022) were expressed under an independent 35S promoter. The images were taken at four days after infiltration of Agrobacteria. The numbers indicate the numbers of leaves that exhibited cell death out of the total numbers. **G)** A non-cell death-inducing RBA (Response to HopBA1) variant (C38A), which lacks 2’,3’-cAMP/cGMP synthetase activity but retains NADase activity (Yu et al., 2022) and PVX resistance activity. Bar plots describe the results of four independent replicates for wild type and three independent replicates for wild type and the *epss* mutant of *N. benthamiana*. Asterisks depict significant differences between treatments as determined by analysis of variance (ANOVA) followed by Tukey’s honest significant difference (HSD) test (*p* < 0.05).

An *Arabidopsis* R protein, RBA1 (Response to HopBA1), which possesses a Toll/ interleukin-1 receptor like domain (TIR domain) induces host cell death upon transient overexpression in *N. benthamiana* (Nishimura et al., 2017, Yu et al., 2022). The TIR domain of RBA1 harbors NADase and nuclease activities, which are responsible for the generation of pRib-AMP/ADP and diADPR/ADPr-ATP and cyclic nucleotide monophosphates (cNMPs) such as 2′,3′-cAMP/cGMP, respectively (Wan et al., 2019; Tian and Li, 2022; Yu et al., 2022). The two enzymatic activities of RBA1 require one of two catalytic residues, namely, Cys83 or Glu86. The RBA1 Cys83, which is a highly conserved among TIR-containing proteins, is responsible for nuclease activity but not for NADase activity. In contrast, RBA1 Glu86 is required for both catalytic activities of RBA1. The NADase activity of RBA1 alone is not sufficient for RBA1 cell death in *N. benthamiana*, because the RBA1(C83A) variant, which lacks nuclease activity, is significantly impaired in cell death induction in *N. benthamiana* upon transient expression (Yu et al., 2022). However, it remains unknown whether the C83A substitution in RBA1, which impairs cell death induction, also impairs disease resistance activity.

To address this question, we used the PVX-YFP-based assay to examine the disease resistance activity of the RBA1(C83A) variant and the RBA1 (E86A) variant as a control. Interestingly, we found that the RBA1 (C83A) variant retained the same disease resistance activity as wild-type RBA1, whereas the RBA1 (E86A) barley showed disease resistance activity to PVX (Fig. 1G). These data suggest that the 2′,3′-cAMP/cGMP synthase activity (i.e., nuclease activity) of RBA1 is required for host cell death but that the remaining NADase activity is sufficient to confer immunity to PVX. Furthermore, the disease resistance activity of the RBA1(C83A) variant was significantly impaired in the *Nbepss* mutant background (Fig. 1G), consistent with the model that TIR NADase products confer immunity via the EDS1 pathway (Huang et al., 2023). In summary, our results have uncoupled the disease resistance activity of RBA1 from the host cell death that is induced by the 2′,3′-cAMP/cGMP synthetase activity. Similar to RBA1, the *Arabidopsis* TIR-containing NLR SNC1 (Suppressor of *npr1-1* constitutive 1) does not seem to require 2′,3′-cAMP/cGMP synthetase activity (i.e., nuclease activity) for disease resistance (Tian et al., 2022). Therefore, it is possible that 2′,3′-cAMP/cGMP produced by TIR enzymes may be sensed by cell death-inducing receptor(s) other than the EDS1 receptor complexes that sense the TIR NADase products. Another possibility is that there may be a threshold for cell death. In this scenario, both TIR NADase and 2′,3′-cAMP/cGMP synthase activities are required to reach the threshold for cell death induction. Finally, we would like to emphasize that disease resistance assays are not intended as a replacement for the widely used cell death assay in the *N. benthamiana* and *N. benthamiana* transient gene expression system; rather, these two assays should be seen as playing complementary roles in illuminating the cell death-independent immunity of R proteins.

## Acknowledgments

We thank Frank Takken, Johannes Stuttmann, Martin Hartmut Schattat, Li Liu, and Alexander Förderer for the pSfinx plasmid and the *Nbeds1 sag101a sag101b pad4* mutant, *RBA1* variant plasmids, and *AvrSr35*/*Sr35* plasmids, respectively. We also thank Neysan Donnelly for editing the manuscript. This work was supported by the Deutsche Forschungsgemeinschaft (DFG, German Research Foundation, SFB-1403–414786233 to J.C., and T.M.). A part of Figure 1 was prepared with BioRender.com.

## Author contributions

K.H., J.C., and T.M. conceived the project. K.H. and T.T. performed the investigations. K.H., T.T., J.C., and T.M. validated the data, J.C. and T.M. supervised the work, K.H., and T.M. wrote the paper with co-author contributions.

## Supplemental Methods

The monomeric YFP coding region was amplified from pXCSG-YFP (Feys et al., 2005) by PCR using primers containing the SfiI recognition sites (SfiIA-mYFPf: TTGGCCATTATGGCCATGGTGAGCAAGGGCGAGGA, SfiIB-mYFPr: TTGGCCGAGGCGGCCTTACTTGTACAGCTCGTCCA), and the PCR product was digested with SfiI (NEB) and ligated into the SfiIA and SfiIB sites of the binary PVX-based expression vector (Takken et al., 2000). The *Agrobacterium tumefaciens* strain GV3101 pMP90RK (Koncz and Schell, 1986) carrying pSfinx-mYFP (OD_600_ = 0.001) and the *A. tumefaciens* strain GV3101 pMP90RK (Koncz and Schell, 1986) carrying resistance genes (OD_600_ = 0.6) were co-infiltrated into leaves of *Nicotiana benthamiana*. At four days after infiltration, ten 5-mm leaf-discs per condition were collected using a biopsy punch from the infiltrated areas. The collected leaf-discs were snap-frozen using liquid nitrogen and kept in a -80 °C freezer until use. The leaf-discs were ground using a pestle or Retsch mill, and 200 μl of the extraction buffer (50 mM Tris-HCl pH 8.0, 150 mM NaCl, proteinase inhibitor (Roche: cOmplet, EDTA-free Protease Inhibitor Cocktail)) were added. The resulting lysate was centrifuged for 5 min at 18,000 x g at 4 °C. The supernatant was transferred into a new tube and centrifuged for 5 min at 18,000 x g at 4 °C. Then, 80 μl of the supernatant were loaded into a black 96 well plate (Corning: CLS3915-100EA). A fluorescence microplate reader (TECAN: Tecan infinite 200 pro) was used to measure YFP intensity. We configured excitation/emission spectra of 516/560 nm with the Tecan infinite 200 pro. While we note that excitation/emission spectra of 500/540 nm can be similarly used in place of 516/560 nm, the excitation/emission spectra of 516/530 nm, which are theoretically optimal for YFP, did not detect the YFP intensity. This may be due to the small Stokes shifts of YFPs. The emission spectra of leaf extracts with expression of PVX-YFP or PVX shown in Fig. 1C were determined by a confocal microscope using the lambda scan function (Leica, STELLARIS 5). The fluorescence stereomicroscopic images of the abaxial leaf surface of *N. benthamiana* shown in Fig. 1D were obtained using a fluorescence stereomicroscope (Leica, MZ16 FA) with a GFP filter equipped with a CCD color sensor. In this assay, we infiltrated the *Agrobacteria* carrying pSfinx-mYFP with OD_600_ = 0.01, 0.001 or 0.0001 and the samples were examined at four days after infiltration. For the Sr35 assay in Fig. 1F, *A. tumefaciens* strain GV3101 pMP90RK carrying Sr35 (OD_600_ = 0.5) and AvrSr35(OD_600_ = 0.075) (Förderer et al., 2022) were co-infiltrated with *A. tumefaciens* carrying pSfinx-mYFP (OD_600_ = 0.001). The images and leaf samples were taken at four days after infiltration of Agrobacteria. For the RBA assay in Fig. 1F, *A. tumefaciens* strain carrying pSfinx-mYFP (OD_600_ = 0.001) and the *A. tumefaciens* strain carrying either of GUS or RBA1 wild-type, C83A, and E86A (OD_600_ = 0.6) were co-infiltrated into leaves of wild type or the *eds1 pad4 sag101a sag101b* mutant (Gantner et al., 2019) of *N. benthamiana* (*Nbepss*).

## Notes

### Competing Interest Statement

The authors have declared no competing interest.

